# Large language models recover archival pathology data and identify NSE as a candidate prognostic marker in Ewing sarcoma

**DOI:** 10.64898/2026.02.20.707103

**Authors:** Jingwei Huang, Ayesha Batool, Zifan Gu, Zhuo Zhao, Bo Yao, Jennifer O. Black, Jessica L. Davis, Alyaa al-Ibraheemi, Steven G. DuBois, Riaz Gillani, Donald A. Barkauskas, Subhash Ramakrishnan, David Hall, Patrick Grohar, Olivier Delattre, Yang Xie, Ling Cai, Guanghua Xiao, Patrick J. Leavey

## Abstract

**Background:** Immunophenotypic information in cooperative-group trial archives remains difficult to analyze. We investigated whether large language models (LLMs) could recover this information from scanned reports and identify prognostic markers in Ewing sarcoma.

**Methods:** An LLM pipeline extracted 18 immunostains from reports collected through six Children’s Oncology Group (COG) studies. Accuracy was evaluated against clinician annotation. Overall survival (OS) associations were screened in three nested populations: the full cohort, cases with molecular support, and fusion-confirmed cases, with false-discovery-rate (FDR) adjustment within each population. The selected marker was evaluated at the transcript level in two independent cohorts.

**Findings:** Of 931 patients from 185 institutions, 928 had evaluable reports. Mean extraction accuracy was 98.2%. Neuron-specific enolase (NSE) was recorded in 164 patients (18%). NSE ranked first in all three screens and met the FDR threshold for OS in the fusion-confirmed subset (adjusted p = 0.008; 0 of 22 NSE-negative versus 11 of 34 NSE-positive patients died). Among 162 patients with NSE and OS data, positivity was associated with inferior OS (5-year OS 61% versus 85%; HR 2.17, 95% CI 1.17–4.05), including after adjustment for metastatic status (HR 2.13, 1.13–3.99). An exploratory analysis suggested a stronger association in localized disease (HR 5.62, 1.66–19.1; interaction p = 0.038). High *ENO2* transcript (encoding NSE) expression was associated with inferior OS in both external cohorts (HR 1.86, 1.04–3.32, and 3.94, 1.65– 9.43).

**Interpretation:** LLM-assisted extraction enables prognostic investigation of archival pathology reports. NSE is a candidate adverse prognostic marker, with independent transcript-level supporting evidence. Standardized protein-level validation is warranted, particularly in localized Ewing sarcoma.

**Research in Context:** *Evidence before this study:* We searched PubMed and Google Scholar for articles published up to October 2026 using the terms “Ewing sarcoma”, “prognosis”, “risk stratification”, “circulating tumor DNA”, “immunohistochemistry”, “large language models”, “natural language processing”, and “pathology report extraction”. Risk stratification in Ewing sarcoma relies on clinical factors including metastatic status, tumor size and age. STAG2 alteration and pretreatment circulating tumor DNA burden have recently been validated as prognostic in Children’s Oncology Group cohorts and incorporated into proposed clinical-molecular risk groups, and a Europe-wide screen of prospectively collected tumors identified MIR34A expression, LOH, and PGA as candidate biomarkers; no immunohistochemical marker has been validated for prognosis. Small studies from the pre-molecular era reported inconsistent associations between neural-differentiation markers, including neuron-specific enolase (NSE), and outcome. Our search did not identify a comparable study combining immunostain screening in a large cooperative-group archive, analyses restricted by molecular diagnostic evidence, and independent transcript-level evaluation. Previous studies support LLM extraction from clinical and pathology text, including scanned reports, but evidence linking extraction from heterogeneous trial archives to biomarker discovery remains limited.

*Added value of this study:* We analyzed an archive of pathology reports from 931 patients enrolled in six COG studies at 185 institutions over 21 years. The LLM pipeline recovered structured immunostain data from noisy OCR text with 98.2% mean accuracy against a clinician-annotated reference. OS screening identified NSE as the top-ranked marker in the full cohort and in two nested populations defined by molecular diagnostic evidence. NSE met the FDR threshold among fusion-confirmed cases. Its association with OS persisted after adjustment for metastatic status and received independent support from *ENO2* transcript (encoding NSE) expression in two cohorts that were not processed by the extraction pipeline.

*Implications of all the available evidence:* Cooperative-group pathology archives can support biomarker discovery when narrative information is recovered, and its extraction accuracy assessed. The NSE/*ENO2* findings support standardized evaluation of NSE as a candidate prognostic marker, including assessment of its contribution beyond clinical factors. The apparent stronger OS association in localized disease warrants further investigation. Our work establishes the use of LLMs as a strategy for learning from vast archives of cooperative group data in other rare cancers.

## Introduction

Ewing sarcoma is an aggressive bone and soft-tissue malignancy of children and young adults. Despite intensive multimodal therapy, outcomes remain heterogeneous, and risk stratification relies principally on clinical features such as metastatic status, tumor size and age.^1–3^ Recent studies^4–8^ have identified and validated molecular prognostic markers, including STAG2 and TP53 alterations and pretreatment circulating tumor DNA burden. Diagnosis integrates morphology, immunohistochemistry and molecular evidence of an FET::ETS fusion.^6,9^ Related round-cell sarcomas with CIC rearrangements or BCOR alterations are classified separately and have distinct clinicopathological features.^10,11^ Histologic and immunophenotypic variation within Ewing sarcoma may also carry prognostic information, but its evaluation across large trial populations has been limited.

One obstacle is the format of archival data. The Children’s Oncology Group (COG) has collected scanned institutional diagnostic pathology reports over more than two decades. These reports document the immunostains individual pathologists ordered and their results, across institutions and changing diagnostic practice. Their heterogeneous layouts and variable image quality make systematic abstraction difficult. LLMs offer a means of extracting structured variables from clinical and pathology reports,^12–15^ provided that performance is assessed on the degraded, optically recognized text found in historical archives.

We developed and evaluated an LLM pipeline to extract immunostain results from pathology reports for 931 patients enrolled in six COG studies. We then screened the recovered variables for association with overall survival (OS) in the full cohort and in nested populations with increasing molecular support for the diagnosis. The resulting NSE candidate was evaluated through *ENO2* transcript (encoding NSE) expression in two independent cohorts. This design connects the recovery of archival pathology information with the investigation of a clinically relevant prognostic hypothesis.

## Methods

### Study design and data sources

We conducted a retrospective cohort study of patients enrolled in six COG studies: the interventional trials AEWS0031 (n = 109), AEWS1031 (n = 84) and AEWS1221 (n = 220), whose primary outcomes have been reported,^1–3^ and the biology studies AEWS02B1 (n = 22), APEC14B1 (n = 312) and AEWS07B1 (n = 184). Institutional diagnosis of Ewing sarcoma was sufficient for eligibility; central pathology review and molecular confirmation were not required for inclusion in this historical archive. COG provided 933 scanned reports corresponding to 931 unique patients, including two duplicate scans, from 165 United States and 20 international institutions. Reports dated from July 2001 to September 2022 were linked to baseline characteristics and survival data supplied by the COG Statistical Data Center (Table 1). After exclusion of two unreadable scans and one non-pathology document, 928 patients had evaluable reports. The study was a secondary analysis of de-identified data approved by the UT Southwestern Institutional Review Board. Figure 1A summarizes extraction and accuracy evaluation; Figure 1B shows the patient populations, and Figure 1C links the immunostain screen to external transcript-level assessment.

**Fig. 1.**
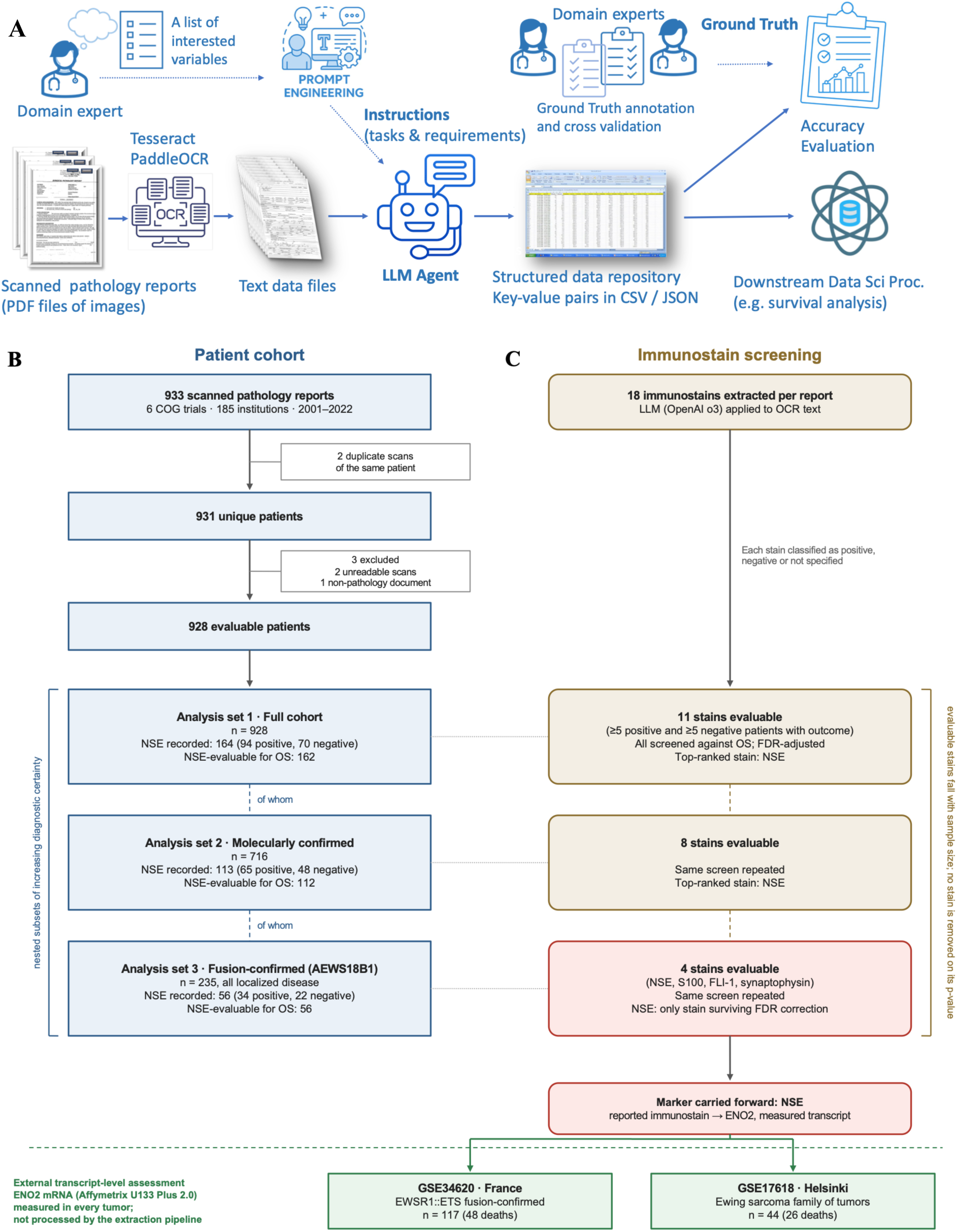
Study overview. (**A**) Scanned institutional pathology reports from six COG studies were converted to text using Tesseract OCR and processed without manual text correction by OpenAI o3 to extract 18 immunostains and documented molecular results. Extraction was evaluated against a 196-report clinician-annotated reference, including 48 independently double-annotated reports assessed against consensus. (**B**) Patient accounting and nested analysis populations: 933 scans represented 931 unique patients after removal of two duplicate scans; exclusion of two unreadable scans and one non-pathology document left 928 evaluable patients. The molecularly supported and fusion-confirmed subsets contained 716 and 235 patients, respectively. NSE was recorded in 164, 113 and 56 patients, of whom 162, 112 and 56 had OS data. (C) Immunostain screening within the corresponding populations yielded 11, eight and four evaluable stains. Evaluability required at least five positive and five negative patients with outcome data; marker counts reflect this criterion within each population, not selection by earlier p-values. NSE was carried forward to independent transcript-level assessment of *ENO2* in GSE34620 (n = 117) and GSE17618 (n = 44). The COG populations overlap; the expression cohorts are independent of the COG archive and extraction pipeline.

**Table 1.** Baseline characteristics of the Ewing sarcoma cohort (N = 931)

| **Characteristic** | **Value** |
|---|---|
| Age at enrollment, years, median (Q1, Q3) | 13.5 (9.5, 16.6) |
| Sex: female / male / unknown | 420 (45%) / 510 (55%) / 1 (0.1%) |
| Race: White | 779 (84%) |
| Asian | 22 (2.4%) |
| Black or African American | 22 (2.4%) |
| Native Hawaiian or other Pacific Islander | 11 (1.2%) |
| American Indian or Alaska Native | 6 (0.6%) |
| Multiple races | 6 (0.6%) |
| Not reported / unknown | 13 (1.4%) / 72 (7.7%) |
| Ethnicity: Hispanic or Latino | 129 (14%) |
| Not Hispanic or Latino | 757 (81%) |
| Not reported / unknown | 6 (0.6%) / 39 (4.2%) |
| Metastatic at diagnosis: yes / no / unknown | 327 (35%) / 597 (64%) / 7 (0.8%) |
| Primary site: axial skeleton | 277 (30%) |
| Long bones | 226 (24%) |
| Soft tissue | 94 (10%) |
| Other / unknown | 287 (31%) / 47 (5.0%) |
| Follow-up, days, median (Q1, Q3) | 1,602 (965, 2,477) |
| Life status: alive / dead / unknown | 657 (70%) / 269 (29%) / 5 (0.5%) |

### Optical character recognition and LLM extraction

Reports varied in scan quality, formatting and language (English and French). Images were converted to text with Tesseract, an open-source optical character recognition (OCR) engine. The raw output, which included misrecognized characters, scan-line artefacts and disrupted layout, was passed to the LLM without manual correction. OpenAI’s o3 model was prompted to classify each of 18 immunostains as Positive, Negative or Not specified: CD99 (MIC2), NSE, S100, NKX2.2, FLI-1, ERG, myogenin, MyoD1, DUX4, WT1, ETV4, BCOR, LCA (CD45), synaptophysin, chromogranin, CCNB3, desmin and vimentin. The panel included stains described in College of American Pathologists (CAP) guidance,^9^ markers reported in the Ewing sarcoma literature, and stains used in the differential diagnosis of round-cell sarcomas. The pipeline also extracted molecular results documented in the reports. The extraction prompt and output schema are provided in Supplementary Figures S1 and S2.

### Accuracy evaluation

A reference standard was created from 200 randomly selected reports; four were excluded (one duplicate, two unreadable scans and one non-pathology document), leaving 196. Two clinicians (AB and PL) annotated the original PDF images. AB annotated all 196 reports, and PL independently reviewed a random subset of 48 reports while blinded to AB’s annotations. Discrepancies in the double-annotated subset were resolved by consensus. Accuracy, precision, recall and F1 were calculated against the reference standard; precision and F1 were weighted by class frequency. In the 48-report subset, the LLM and each annotator were compared with the consensus reference. Table 2 summarizes performance across the reference reports, including the “Not specified” category. Table 3 reports accuracy among reports in which a stain was specified by at least one source, weighted across stains by the number of such reports.

**Table 2.**
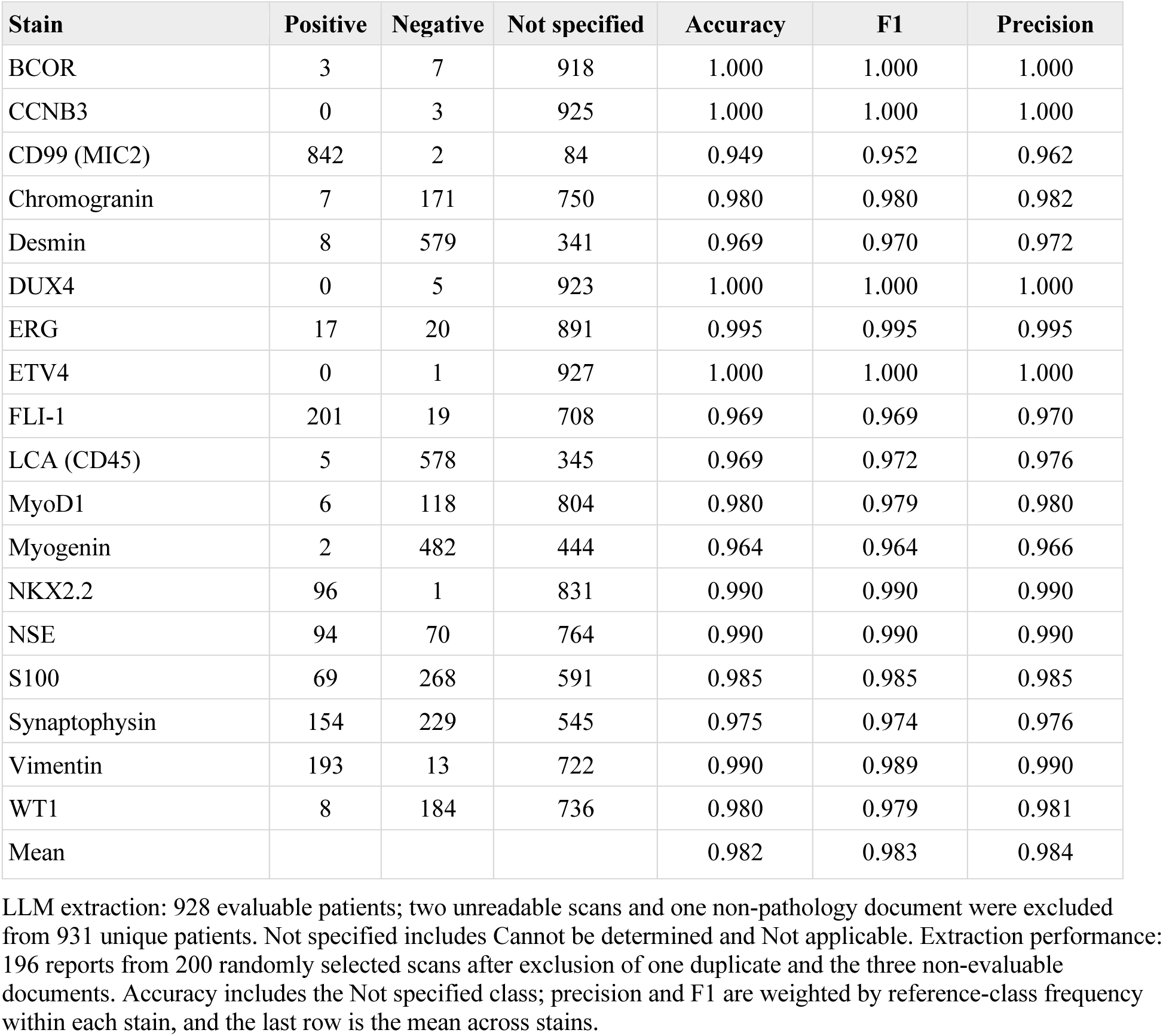
Immunostain results extracted by the LLM and extraction accuracy.

**Table 3.**
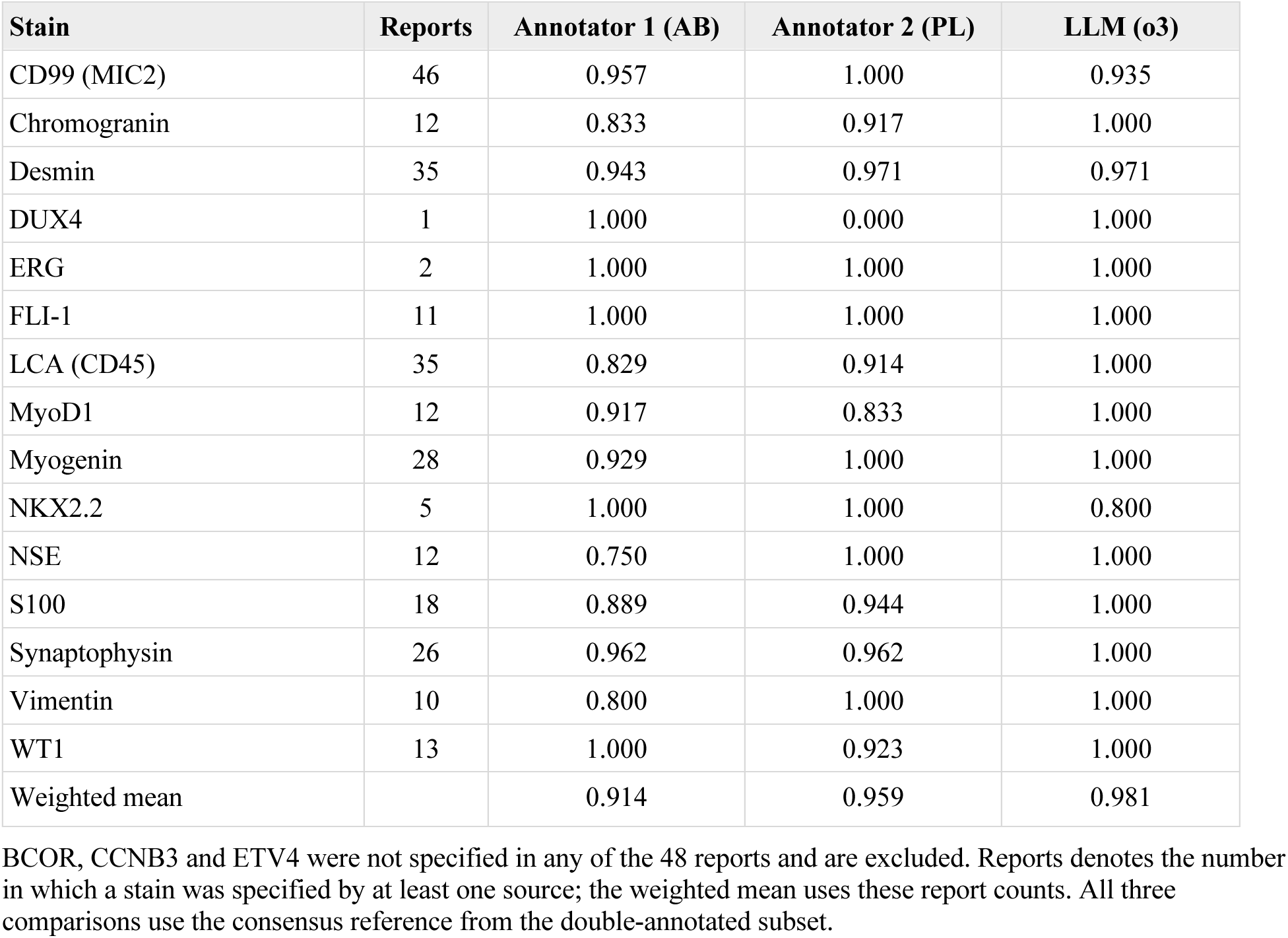
Accuracy of the LLM and two human annotators against the consensus reference in the 48-report blinded subset.

| **Stain** | **Reports** | **Annotator 1 (AB)** | **Annotator 2 (PL)** | **LLM (o3)** |
|---|---|---|---|---|
| CD99 (MIC2) | 46 | 0.957 | 1.000 | 0.935 |
| Chromogranin | 12 | 0.833 | 0.917 | 1.000 |
| Desmin | 35 | 0.943 | 0.971 | 0.971 |
| DUX4 | 1 | 1.000 | 0.000 | 1.000 |
| ERG | 2 | 1.000 | 1.000 | 1.000 |
| FLI-1 | 11 | 1.000 | 1.000 | 1.000 |
| LCA (CD45) | 35 | 0.829 | 0.914 | 1.000 |
| MyoD1 | 12 | 0.917 | 0.833 | 1.000 |
| Myogenin | 28 | 0.929 | 1.000 | 1.000 |
| NKX2.2 | 5 | 1.000 | 1.000 | 0.800 |
| NSE | 12 | 0.750 | 1.000 | 1.000 |
| S100 | 18 | 0.889 | 0.944 | 1.000 |
| Synaptophysin | 26 | 0.962 | 0.962 | 1.000 |
| Vimentin | 10 | 0.800 | 1.000 | 1.000 |
| WT1 | 13 | 1.000 | 0.923 | 1.000 |
| Weighted mean |  | 0.914 | 0.959 | 0.981 |

BCOR, CCNB3 and ETV4 were not specified in any of the 48 reports and are excluded. Reports denotes the number in which a stain was specified by at least one source; the weighted mean uses these report counts. All three comparisons use the consensus reference from the double-annotated subset.

### Analysis populations and endpoints

OS, measured from diagnosis to death from any cause, was the primary endpoint. Event-free survival (EFS), measured from diagnosis to relapse, progression, second malignancy or death, was secondary; detailed EFS results are reported in the supplement. Both endpoints were taken from the COG dataset. Immunostains were screened in three nested populations (Figure 1B–C). The full cohort included all 928 patients with evaluable reports. The molecularly supported subset (n = 716) included patients with an EWSR1 rearrangement documented in AEWS1221 testing^16^, an *EWSR1::FLI1*, *EWSR1::ERG*, *EWSR1::ETV1* or *EWSR1::FEV* fusion identified in the published dataset from the AEWS18B1-Q study^7^ (hereafter AEWS18B1), or a *FET::ETS* fusion or EWSR1 rearrangement recorded in the pathology report. The fusion-confirmed subset comprised 235 AEWS18B1 patients with one of these four EWSR1::ETS fusions. The broader molecularly supported definition includes rearrangements without an identified fusion partner. The subsets share patients and assess the robustness of associations to diagnostic restriction. Within each population, a stain was evaluable if at least five positive and five negative patients had outcome data. This criterion yielded eleven, eight and four evaluable stains, respectively; markers were not removed from subsequent screens on the basis of their earlier p-values. NSE was selected for independent transcript-level evaluation on the basis of the OS screening results.

### External cohorts

Expression of *ENO2*, the gene encoding NSE, was analyzed in two independent Ewing sarcoma cohorts profiled on the Affymetrix HG-U133 Plus 2.0 array. GSE34620 comprises 117 tumors from France with documented EWSR1::ETS fusions;^17^ survival data were provided by the original investigators. GSE17618 comprises 44 tumors of the Ewing sarcoma family from Helsinki;^18^ clinical outcomes were obtained from the source study. Per-sample fusion status is not publicly available for GSE17618. All 44 tumors were included, although the cohort contains some treated and recurrent specimens. Normalized log2 expression values were obtained from the Gene Expression Omnibus. *ENO2* expression at or above the cohort median was classified as high, and expression below the median as low. Tertile analyses were used to examine the pattern of association. This assessment used a different assay from NSE immunohistochemistry and did not involve the OCR–LLM pipeline.

### Statistical analysis

Survival was estimated by the Kaplan–Meier method^19^ and compared using two-sided log-rank tests. Hazard ratios (HR) and 95% confidence intervals (CI) were estimated by Cox regression.^20^ Positive and negative immunostain results were analyzed as a binary variable; Not specified results were treated as missing. For the marker screens, Benjamini–Hochberg adjustment was applied to the log-rank p-values across the stains evaluable within each population and endpoint. An adjusted p < 0.05 was the false discovery rate (FDR) threshold. A sensitivity calculation applied an 11-test Bonferroni penalty to the fusion-subset NSE OS p-value. In the full cohort, multivariable models included NSE and metastatic status at diagnosis. Subgroup analyses by metastatic status were accompanied by an NSE × metastatic-status interaction test; the interaction estimate is reported as the ratio of the NSE HR in localized disease to that in metastatic disease. Adjusted, subgroup and interaction-model p-values were Wald tests unless identified otherwise. When one group had no events, Firth penalized Cox estimates with profile-likelihood confidence intervals and tests were used (Supplementary Table S1). External models were adjusted for age in GSE34620, the only covariate available, and metastatic status in GSE17618. Proportional hazards were assessed using Schoenfeld residuals; external sensitivity analyses included restriction to 60 months and time-split models at 24 months. Targeted external and clinical subgroup analyses are reported with nominal p-values. Analyses used R with the survival, survminer and coxphf packages.

## Results

### Extraction accuracy

The pipeline processed reports from 928 of 931 unique patients (Figure 1A–B). Across 18 stains, 73% of the 16,704 stain–patient combinations were classified as “Not specified” (Table 2). Results were recorded for more than half of patients for four stains: CD99, desmin, LCA and myogenin. NSE was recorded for 164 patients (18%; 94 positive and 70 negative). Against the 196-report reference standard, mean accuracy across the 18 stains was 0.982, with mean weighted precision of 0.984 and mean weighted F1 of 0.983 (Table 2). Figure 2 illustrates extraction performance and the effects of degraded OCR text. In the 48-report double-annotated subset, agreement between the human annotators averaged 96.3%. Against the consensus reference, weighted accuracy was 98.1% for the LLM, 91.4% for AB and 95.9% for PL across the 15 stains represented (Table 3). These descriptive comparisons were based on a small subset. Extraction errors included failure to recover results from degraded OCR text, highlighting the importance of source-image review when automated extraction is uncertain.

**Fig. 2.**
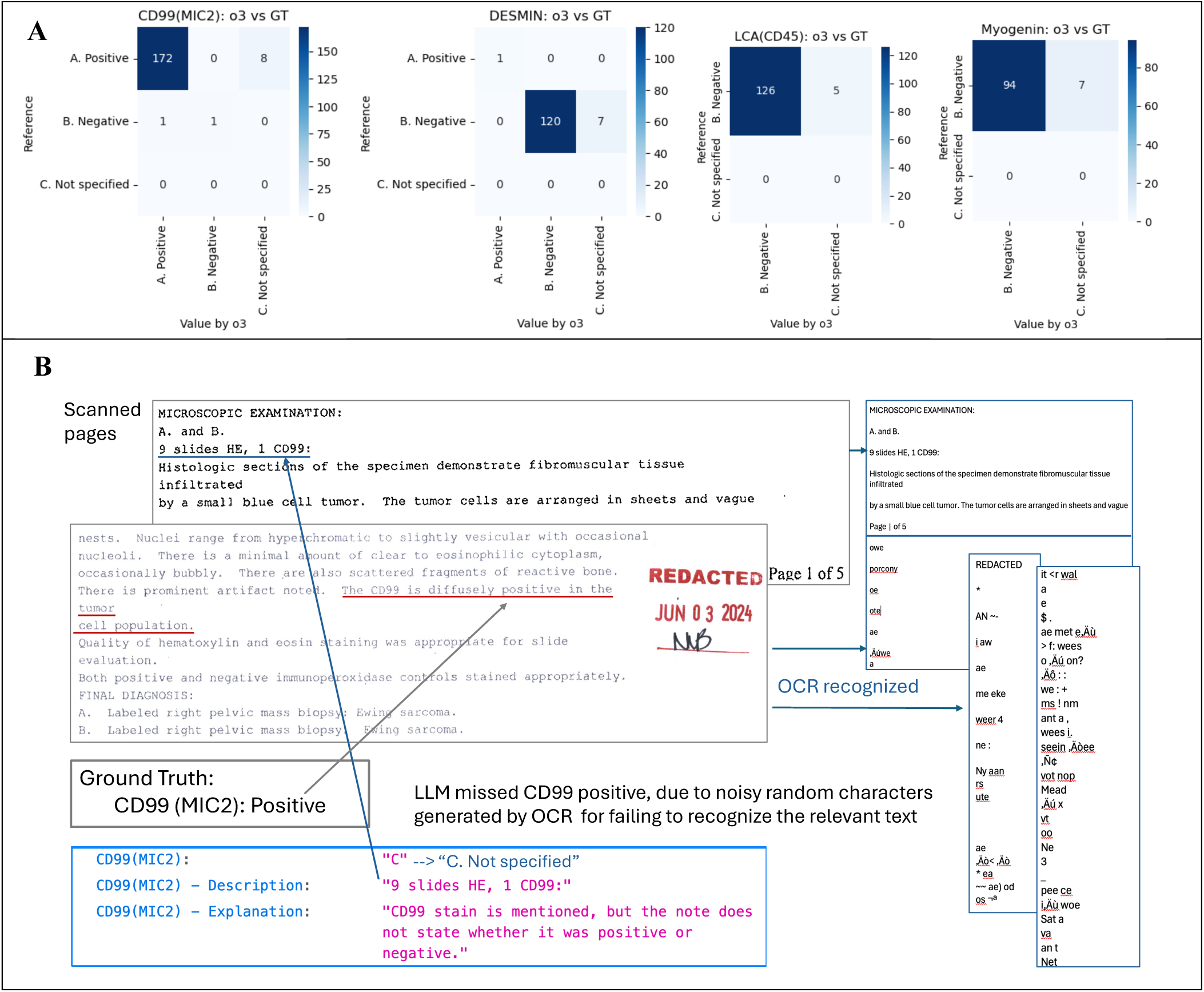
Extraction accuracy. (**A**) Confusion matrices for CD99, desmin, NSE and S100 against the 196-report reference standard, with Positive, Negative and Not specified classes. (**B**) A representative scanned report and its OCR output illustrate character insertion and misalignment that can impair extraction. Table 2 reports summary extraction performance.

### Overall survival screening across three diagnostic populations

The same OS screen was applied within each population (Figure 1C and Table 4). In the full cohort, 11 stains were evaluable. CD99 and NKX2.2 were not evaluable because almost all tumors with recorded staining were positive; myogenin, BCOR, CCNB3, DUX4 and ETV4 had fewer than five positive tumors. Three stains had a raw p-value < 0.05: NSE (HR 2.17, 95% CI 1.17–4.05; log-rank p = 0.012), S100 (HR 0.56, 0.33–0.95; p = 0.029) and desmin (HR 2.80, 1.04–7.55; p = 0.034). None met the Benjamini-Hochberg threshold of adjusted p < 0.05 across the 11 evaluable stains. NSE had the smallest unadjusted p-value (log-rank p = 0.012); after adjustment it was tied with S100 and desmin at p = 0.12. Of the 164 patients with NSE recorded, 162 had OS data (93 positive, with 35 deaths; 69 negative, with 14 deaths). Five-year OS was 60.5% (95% CI 50.4–72.7) in NSE-positive patients and 84.6% (76.3–93.9) in NSE-negative patients (Figure 3A).

**Fig. 3.**
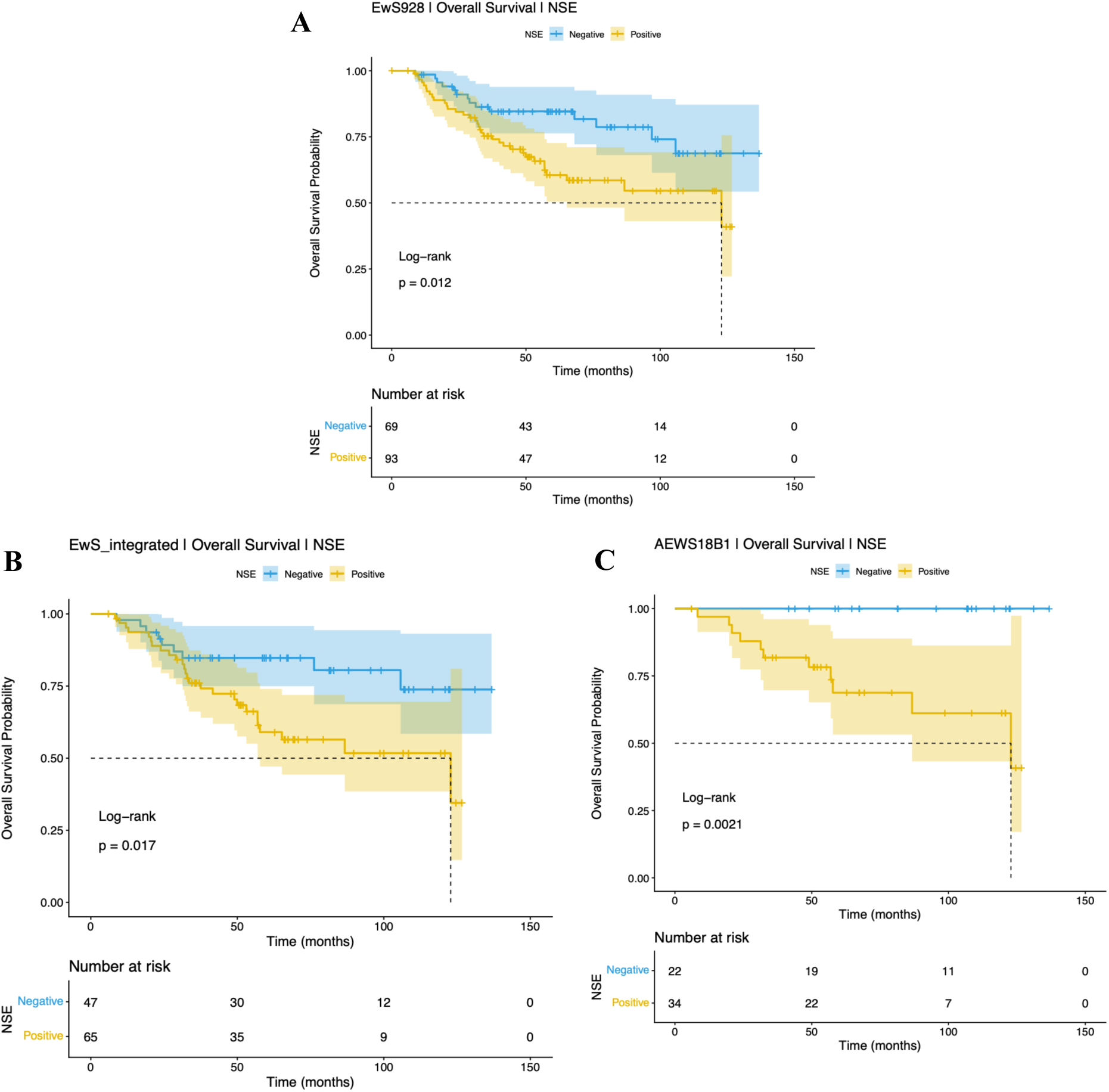
Overall survival by NSE status in three nested populations. Kaplan–Meier curves for OS in (**A**) the full cohort with NSE and OS data (n = 162), (**B**) the molecularly supported subset with NSE and OS data (n = 112), and (**C**) the fusion-confirmed subset (n = 56). Shaded bands indicate 95% confidence intervals; tick marks indicate censoring; numbers at risk are shown below each panel. Time is measured in months from diagnosis. Displayed p-values are log-rank tests. Table 4 reports marker-specific sample sizes, Cox HRs and BH-adjusted p-values for each screen. No NSE-negative patient died in the fusion-confirmed subset; the Firth penalized estimate is reported in Supplementary Table S1. The three populations overlap and are not independent validation cohorts.

**Table 4.** Overall survival screen across three nested diagnostic populations.

|  | Full cohort (n = 928) — 11 evaluable stains |  |  |  | Molecularly supported (n = 716) — 8 evaluable stains |  |  |  | Fusion-confirmed, AEWS18B1 (n = 235) — 4 evaluable stains |  |  |  |
|---|---|---|---|---|---|---|---|---|---|---|---|---|
| Stain | Neg / Pos n (events) | HR (95% CI) | Log-rank p | BH adj. p | Neg / Pos n (events) | HR (95% CI) | Log-rank p | BH adj. p | Neg / Pos n (events) | HR (95% CI) | Log-rank p | BH adj. p |
| **NSE** | **69 (14) / 93 (35)** | **2.17 (1.17–4.05)** | **0.012** | **0.12** | **47 (9) / 65 (26)** | **2.46 (1.15–5.27)** | **0.017** | **0.13** | **22 (0) / 34 (11)** | **NE‡** | **0.002** | **0.008** |
| S100 | 268 (97) / 69 (16) | 0.56 (0.33–0.95) | 0.029 | 0.12 | 203 (69) / 51 (15) | 0.76 (0.44–1.33) | 0.34 | 0.68 | 59 (11) / 15 (2) | 0.67 (0.15–3.04) | 0.61 | 0.61 |
| Desmin | 576 (180) / 8 (4) | 2.80 (1.04–7.55) | 0.034 | 0.12 | — | not evaluable† |  |  | — | not evaluable† |  |  |
| LCA (CD45) | 574 (174) / 5 (3) | 2.16 (0.69–6.78) | 0.17 | 0.48 | 439 (132) / 5 (3) | 2.25 (0.72–7.08) | 0.15 | 0.61 | — | not evaluable† |  |  |
| FLI-1 | 19 (8) / 201 (60) | 0.68 (0.32–1.42) | 0.30 | 0.66 | 13 (5) / 163 (47) | 0.77 (0.31–1.93) | 0.58 | 0.68 | 5 (2) / 55 (11) | 0.55 (0.12–2.47) | 0.42 | 0.61 |
| Chromogranin | 171 (49) / 7 (1) | 0.47 (0.06–3.37) | 0.44 | 0.71 | — | not evaluable† |  |  | — | not evaluable† |  |  |
| MyoD1 | 117 (36) / 6 (1) | 0.47 (0.06–3.45) | 0.45 | 0.71 | — | not evaluable† |  |  | — | not evaluable† |  |  |
| ERG | 20 (3) / 17 (4) | 1.55 (0.34–6.96) | 0.57 | 0.78 | 15 (2) / 13 (3) | 1.98 (0.33–11.84) | 0.45 | 0.68 | — | not evaluable† |  |  |
| Vimentin | 13 (4) / 191 (52) | 0.80 (0.29–2.22) | 0.67 | 0.82 | 10 (2) / 140 (40) | 1.47 (0.36–6.10) | 0.59 | 0.68 | — | not evaluable† |  |  |
| WT1 | 184 (64) / 8 (3) | 1.19 (0.37–3.78) | 0.77 | 0.85 | 142 (47) / 5 (1) | 0.57 (0.08–4.15) | 0.57 | 0.68 | — | not evaluable† |  |  |
| Synaptophysin | 228 (73) / 154 (48) | 1.00 (0.70–1.44) | 1.00 | 1.00 | 169 (55) / 119 (36) | 0.92 (0.61–1.41) | 0.71 | 0.71 | 55 (9) / 39 (8) | 1.39 (0.54–3.62) | 0.49 | 0.61 |

Positive versus negative staining in three nested populations. HRs and 95% CIs are from univariable Cox models; p-values are log-rank tests, and BH-adjusted p-values control FDR across the stains evaluable within each population. Evaluability required at least five positive and five negative patients with outcome data. Patient counts and deaths are shown for each stain. NSE had the smallest p-value in each population and met the FDR threshold in the fusion-confirmed subset. An 11-test Bonferroni sensitivity calculation for its fusion-subset OS p-value gives approximately 0.023. † Not evaluable: fewer than five patients in a staining group. CD99, NKX2.2, myogenin, BCOR, CCNB3, DUX4 and ETV4 were not evaluable in any population and are omitted. ‡ Standard Cox HR not estimable because no NSE-negative patient died (0 of 22 versus 11 of 34 deaths); the Firth penalized estimate is reported in Supplementary Table S1. The population sizes in the column headings are total cohort sizes; each marker analysis includes only patients with recorded staining and OS data.

In the molecularly supported subset, eight stains were evaluable. NSE was recorded in 113 patients, of whom 112 had OS data (65 positive and 47 negative). NSE again had the smallest p-value (HR 2.46, 95% CI 1.15–5.27; log-rank p = 0.017; adjusted p = 0.13; Figure 3B). The S100 estimate was closer to the null (HR 0.76, 0.44–1.33; p = 0.34), and desmin was no longer evaluable. In the fusion-confirmed subset, four stains were evaluable: NSE, S100, FLI-1 and synaptophysin. All 56 patients with NSE recorded had OS data and localized disease. None of 22 NSE-negative patients died, compared with 11 of 34 NSE-positive patients (5-year OS 100% versus 68.7%, 95% CI 53.2–88.9 for the positive group; log-rank p = 0.002; Figure 3C). NSE was the only stain to meet the FDR threshold (adjusted p = 0.008); the 11-test Bonferroni sensitivity calculation also remained below 0.05 (approximately 0.023). The other three stains did not show statistically significant associations. Firth estimation was required for the NSE OS effect in this subset because the negative group had no deaths; its wide confidence interval limits interpretation of the magnitude (Supplementary Table S1). NSE was carried forward for external transcript-level assessment because it ranked first in all three OS screens and met the FDR threshold in fusion-confirmed disease. EFS analyses provided supporting evidence and are reported in Supplementary Figure S3 and Table S3.

### Independent transcript-level assessment of *ENO2*

We evaluated *ENO2* expression in two independent cohorts in which expression was measured in every included tumor, avoiding discretionary NSE staining and the extraction pipeline used for the COG archive (Figure 4). In GSE34620 (117 patients, 48 deaths), *ENO2*-high tumors had inferior OS (5-year OS 53% versus 78%; HR 1.86, 95% CI 1.04–3.32; log-rank p = 0.033). In GSE17618 (44 patients, 26 deaths), *ENO2*-high tumors were also associated with inferior OS (5-year OS 27% versus 72%; HR 3.94, 1.65–9.43; log-rank p = 0.0009). Adjustment changed the effect estimates little: the age-adjusted HR in GSE34620 was 1.80 (1.00–3.26; Wald p = 0.051), and the metastatic-status-adjusted HR in GSE17618 was 4.88 (1.96–12.2). EFS associations were directionally concordant (Supplementary Figure S4 and Table S2). Tertile analyses showed the largest estimated risks in the highest expression group (Supplementary Table S4). Exploratory cutoff analyses were consistent from the median to approximately the 70th percentile. These patterns support an association between higher ENO2 expression and inferior survival but do not establish a discrete biological threshold, as the hazard ratios depend on the chosen cut. Proportional hazards were not rejected for the OS models, but were violated for EFS in GSE34620, and a 24-month split suggested a larger early effect on OS in that cohort (Supplementary Table S5); the HRs in the external cohorts are therefore best read as associations over follow-up rather than constant effects.

**Fig. 4.**
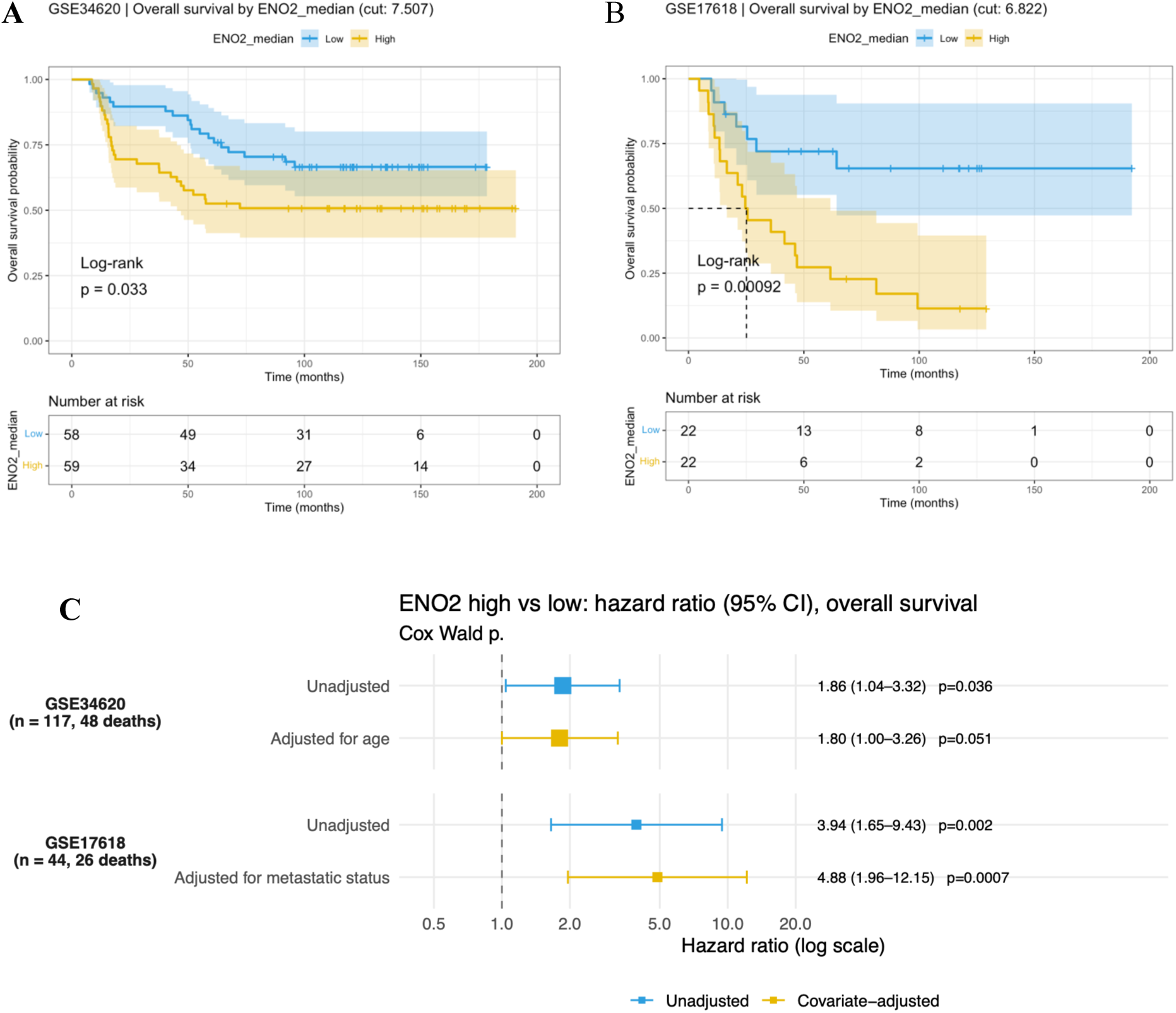
Independent transcript-level assessment of *ENO2* and overall survival. Kaplan–Meier OS curves by *ENO2* expression in (**A**) GSE34620 (n = 117) and (**B)** GSE17618 (n = 44). Expression at or above the cohort median is classified as high, and expression below the median as low. Shaded bands indicate 95% confidence intervals; tick marks indicate censoring; numbers at risk are shown below the curves, and p-values are log-rank tests. (**C**) Unadjusted and covariate-adjusted Cox HRs for high versus low expression, with 95% confidence intervals (adjustment for age in GSE34620 and metastatic status in GSE17618). EFS results are shown in Supplementary Figure S4 and Table S2. Neither cohort was processed by the extraction pipeline.

### Exploratory analyses by metastatic status

The OS analysis with known metastatic status included 160 patients. After adjustment for metastatic status, the NSE HR was 2.13 (95% CI 1.13–3.99; Wald p = 0.019). Among 120 patients with localized disease, NSE positivity was associated with inferior OS (5-year OS 70.5% versus 97.7%; HR 5.62, 1.66–19.1; log-rank p = 0.0018; Figure 5A). Among 40 patients with metastatic disease, the estimate was smaller and less precise (HR 1.22, 0.54–2.78; Wald p = 0.63; Figure 5B). The OS interaction suggested a stronger association in localized disease (ratio of HRs 4.65, 1.09–19.9; Wald p = 0.038). A similar pattern was observed in the molecularly supported subset, where no NSE-negative patient with localized disease died (Supplementary Table S1B). The fusion-confirmed NSE subgroup comprised patients with localized disease only. These overlapping analyses support further investigation in localized disease, but the small event counts and wide confidence intervals limit inference about the size of the interaction. No corresponding interaction was detected for EFS in the full cohort (p = 0.45; Supplementary Table S1B).

**Fig. 5.**
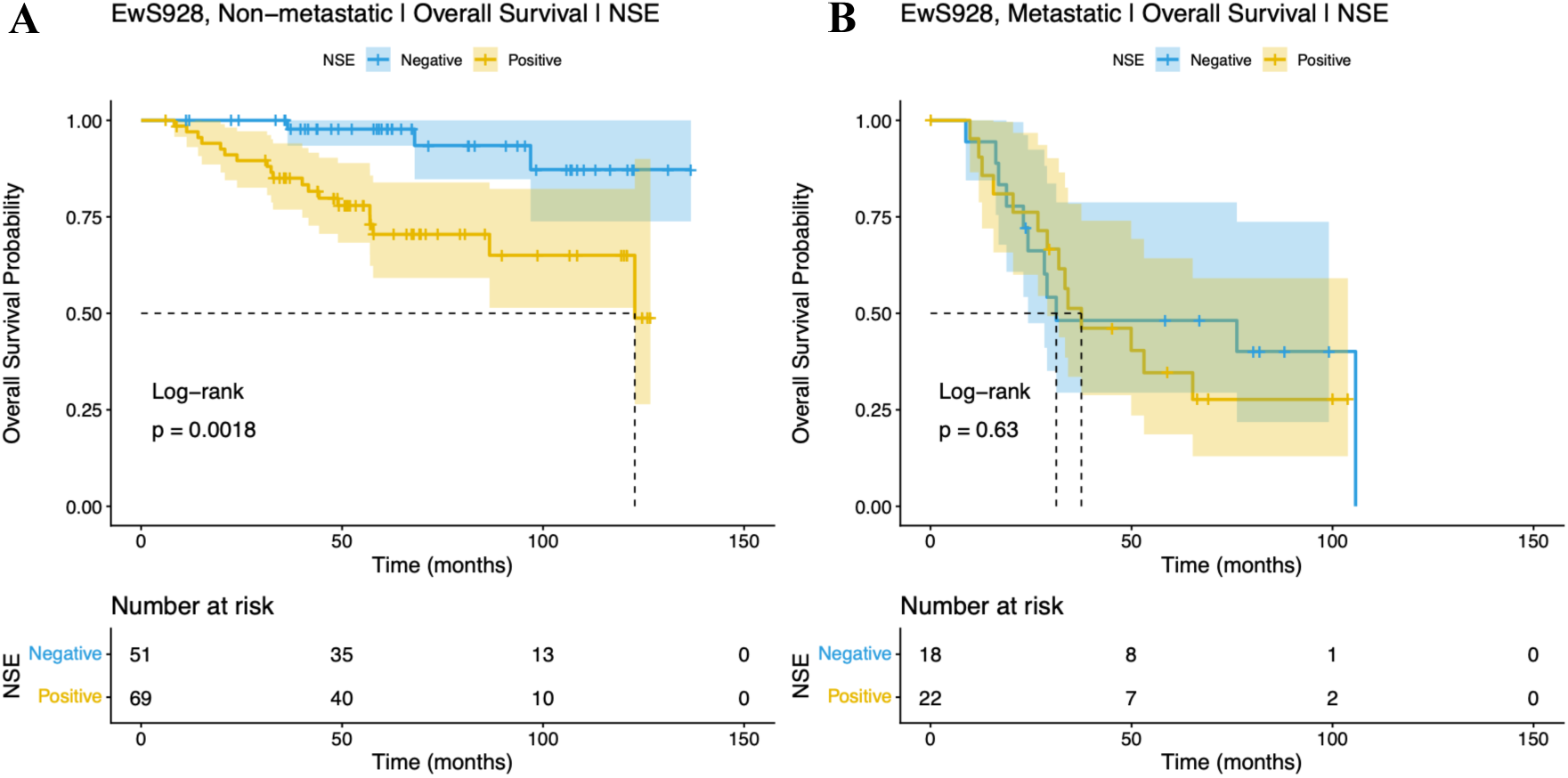
Exploratory analysis of NSE and overall survival by metastatic status. Kaplan–Meier OS curves among patients with (**A**) localized disease (n = 120) and (**B**) metastatic disease (n = 40) in the full cohort. Shaded bands indicate 95% confidence intervals; tick marks indicate censoring; numbers at risk are shown below the curves. Curve-comparison p-values are log-rank tests. Model results include the NSE HR within each subgroup, the NSE HR adjusted for metastatic status, and the NSE × metastatic-status interaction, expressed as the ratio of the NSE HR in localized disease to that in metastatic disease. Adjusted, subgroup and interaction-model p-values are Wald tests. Time is measured in months from diagnosis.

## Discussion

This study connects the recovery of information from archival pathology reports with the discovery of a candidate prognostic marker in Ewing sarcoma. An LLM pipeline extracted immunostain results from evaluable reports for 928 patients in a 931-patient COG archive, with high accuracy against a clinician-annotated reference. NSE had the smallest OS p-value in each of three nested populations and met the FDR threshold in the fusion-confirmed subset. Its association with OS persisted after adjustment for metastatic status, and *ENO2* expression was associated with inferior OS in two independent cohorts. Together, these findings support further evaluation of NSE as a candidate adverse prognostic marker.

NSE is a marker of neural differentiation, providing a biological context for the observed association. Earlier studies examined neural-differentiation markers and outcome in Ewing sarcoma, with inconsistent findings,^21–24^ while subsequent work supported common classification and treatment approaches for tumors historically termed Ewing sarcoma or peripheral primitive neuroectodermal tumor.^24^ The present results suggest that variation in NSE/*ENO2* may retain prognostic relevance within Ewing sarcoma. The exploratory OS analysis points to localized disease as a population for further study, because current clinical categories do not fully explain outcome variation. Whether NSE expression relates to *STAG2* or *TP53* alterations,^4–7^ to neural differentiation, or to transcriptional states associated with the fusion oncoprotein remains to be established. Higher estimated risk in the upper *ENO2* tertile is compatible with several biological models and does not demonstrate a distinct threshold-defined phenotype.

The extraction approach is also relevant to other cooperative-group archives. Previous studies have demonstrated structured information extraction from clinical and pathology text, including bilingual reports, scanned documents and pediatric clinical reports.^12–15,25^ Our study evaluated heterogeneous scans from 185 institutions, two languages and 21 years of diagnostic practice, and linked the recovered variables to trial outcomes. The workflow in Figure 1A provides a basis for evaluating additional report-derived variables. Its use in other diseases or archives will require task-specific validation. The present accuracy estimates include a large Not specified class, and most reference reports were annotated by one clinician; the 48-report consensus subset provides a limited comparison with human performance. These results support the feasibility of automated abstraction but do not establish superiority over expert review.

Several limitations affect interpretation of the biomarker results. NSE was recorded in only 18% of evaluable patients and was ordered at pathologist discretion. The tested subgroup may differ by clinical presentation, institution, era or other characteristics, and the reported adjusted model includes metastatic status alone. Residual confounding by age, tumor site, size or treatment therefore remains possible. Restriction by molecular evidence reduces concern about diagnostic heterogeneity, but the nested analyses are not independent replications. The fusion-confirmed NSE subgroup is small, entirely localized and has no deaths in the negative group; its stronger separation cannot be attributed solely to greater diagnostic certainty. The external expression cohorts provide independent support without discretionary immunostaining, but they do not resolve selection bias in the COG subgroup or validate the NSE IHC assay. *ENO2* mRNA and NSE staining are different measurements; GSE17618 includes some treated and recurrent specimens, and hazards were not proportional in all external models. Independent assessment using pretreatment tissue, standardized staining and scoring, and appropriate clinical adjustment is needed.

LLM-assisted extraction made pathology information from a cooperative-group trial archive available for prognostic investigation. The NSE association in fusion-confirmed disease, together with independent *ENO2* findings, supports standardized protein-level validation and evaluation of added prognostic information beyond clinical factors, particularly in localized Ewing sarcoma. Prospective evaluation with standardized staining and scoring in a molecularly confirmed cohort is planned as part of work toward multimodal prognostic modeling.

## Ethics statement and patient consent

This report describes a secondary analysis of existing, de-identified datasets from the NCI-sponsored Children’s Oncology Group. All patients provided consent for use of collected biospecimens at enrollment. The study involved no interaction with individuals, no access to direct identifiers or re-identification keys, and no new data or specimen collection. The project was approved by the UT Southwestern IRB.

## Acknowledgements

NIH NCI (1R21CA317725, 4R01CA285336), NCTN Operations Center Grant (U10CA180886), NCTN Statistics & Data Center Grant (U10CA180899), COG Biospecimen Bank Grant (U24CA196173); St. Baldrick’s Foundation, Wipe Out Kids’ Cancer Foundation, Quad W Foundation, American Society of Clinical Oncology (ASCO) Career Development Award, Department of Defense Peer-Reviewed Cancer Research Program Fellow Career Development Award, CA220721, NIH K08CA276701, and the Cancer Prevention and Research Institute of Texas (RP25056, RP240521). The content is solely the responsibility of the authors and does not necessarily represent the official views of the National Institutes of Health.

## Supplementary material (list)

**Figure S1.**
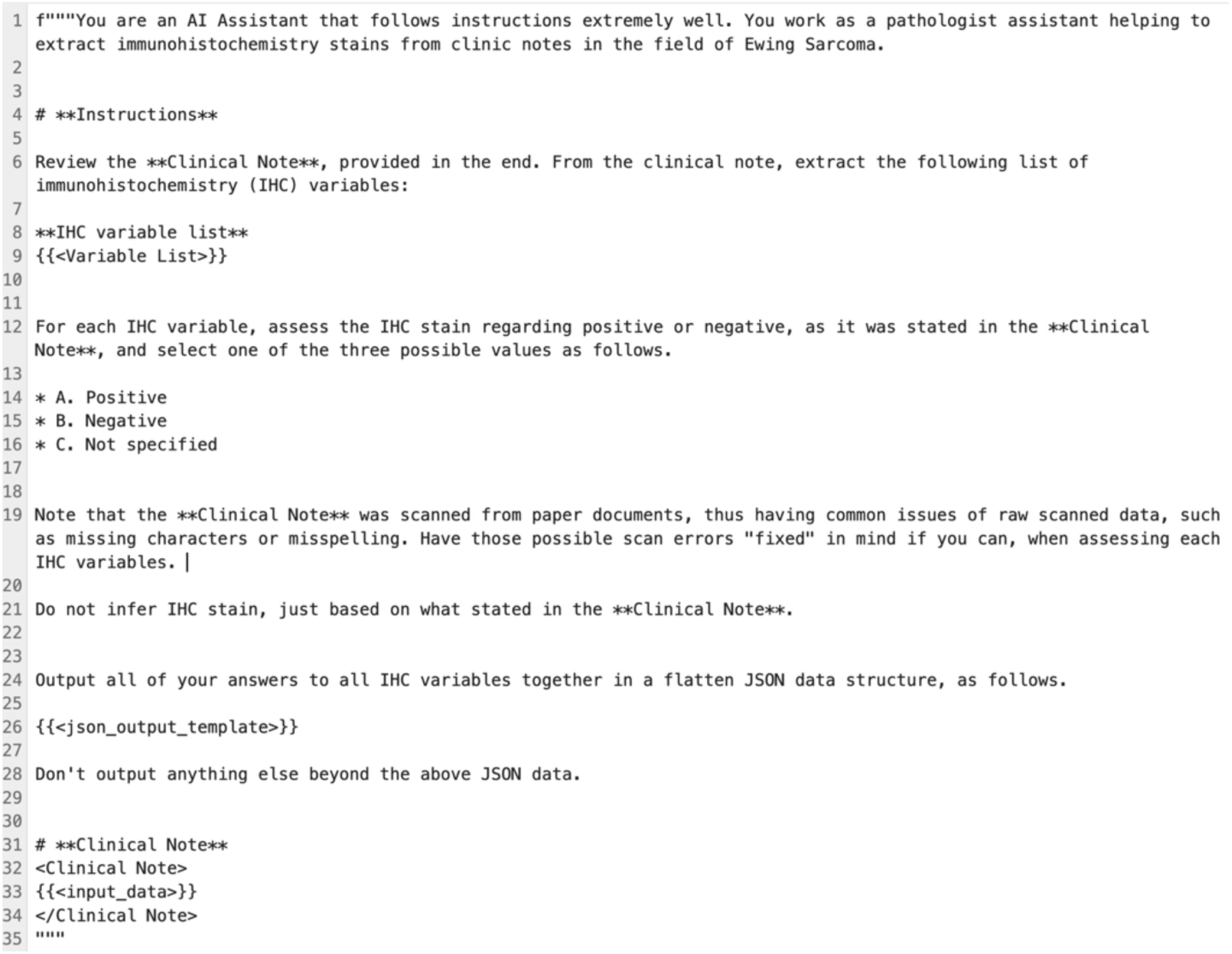
Prompt for immunostain extraction. Prompt template for IHC data extraction. In the template, placeholder {{<Variable List>}} will be replaced with CD99(MIC2), NKX2.2, FLI-1, ERG, Myogenin, MyoD1, DUX4, WT1, ETV4, BCOR, LCA(CD45), SYNAPTOPHYSIN, CHROMOGRANIN, DESMIN, and VIMENTIN. {{<json_output_template>}} will be replaced with the JSON output template provided. {{<input_data>}} will be replaced with each input pathology report.

**Figure S2.**
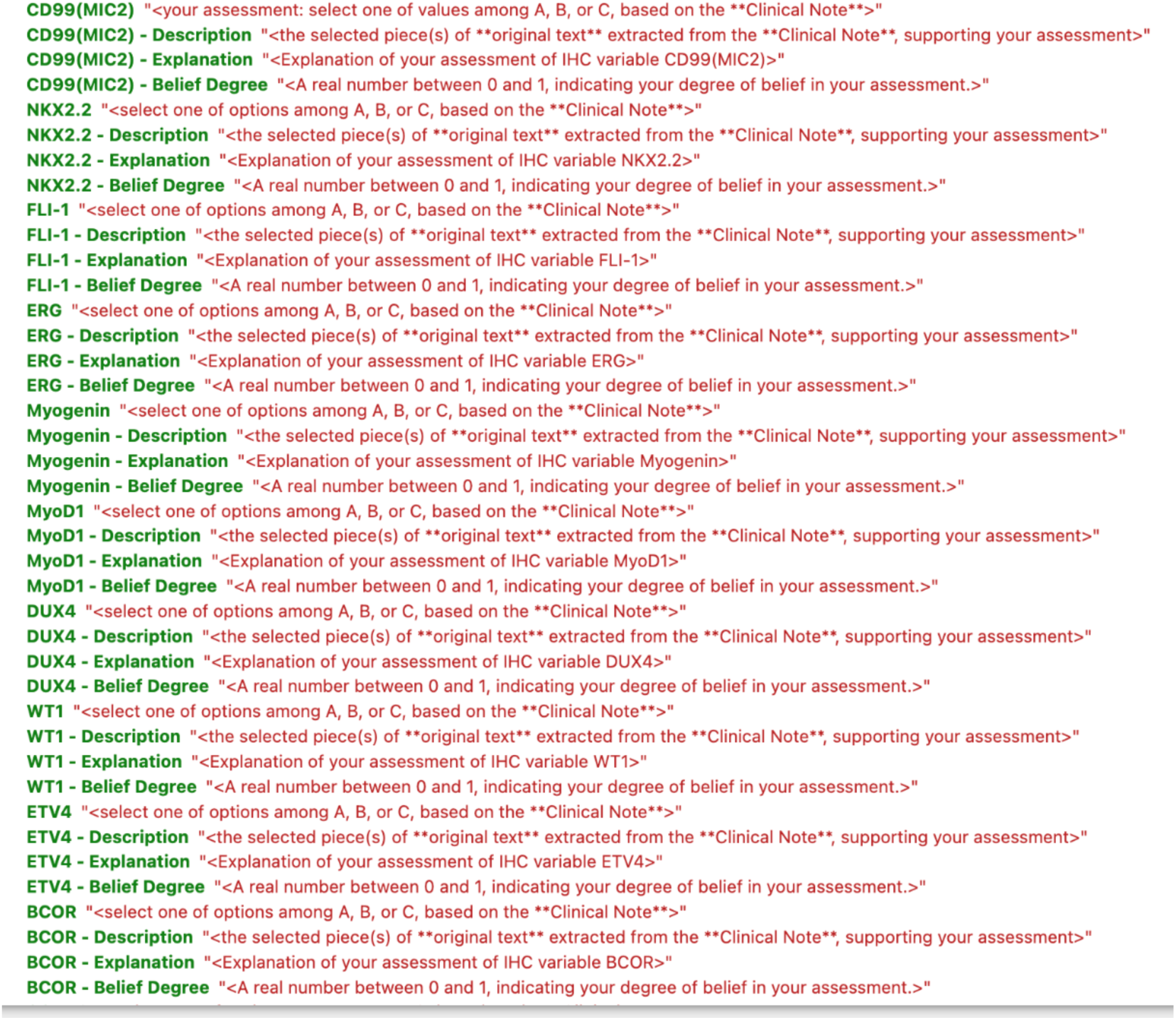
JSON output template.

**Figure S3.**
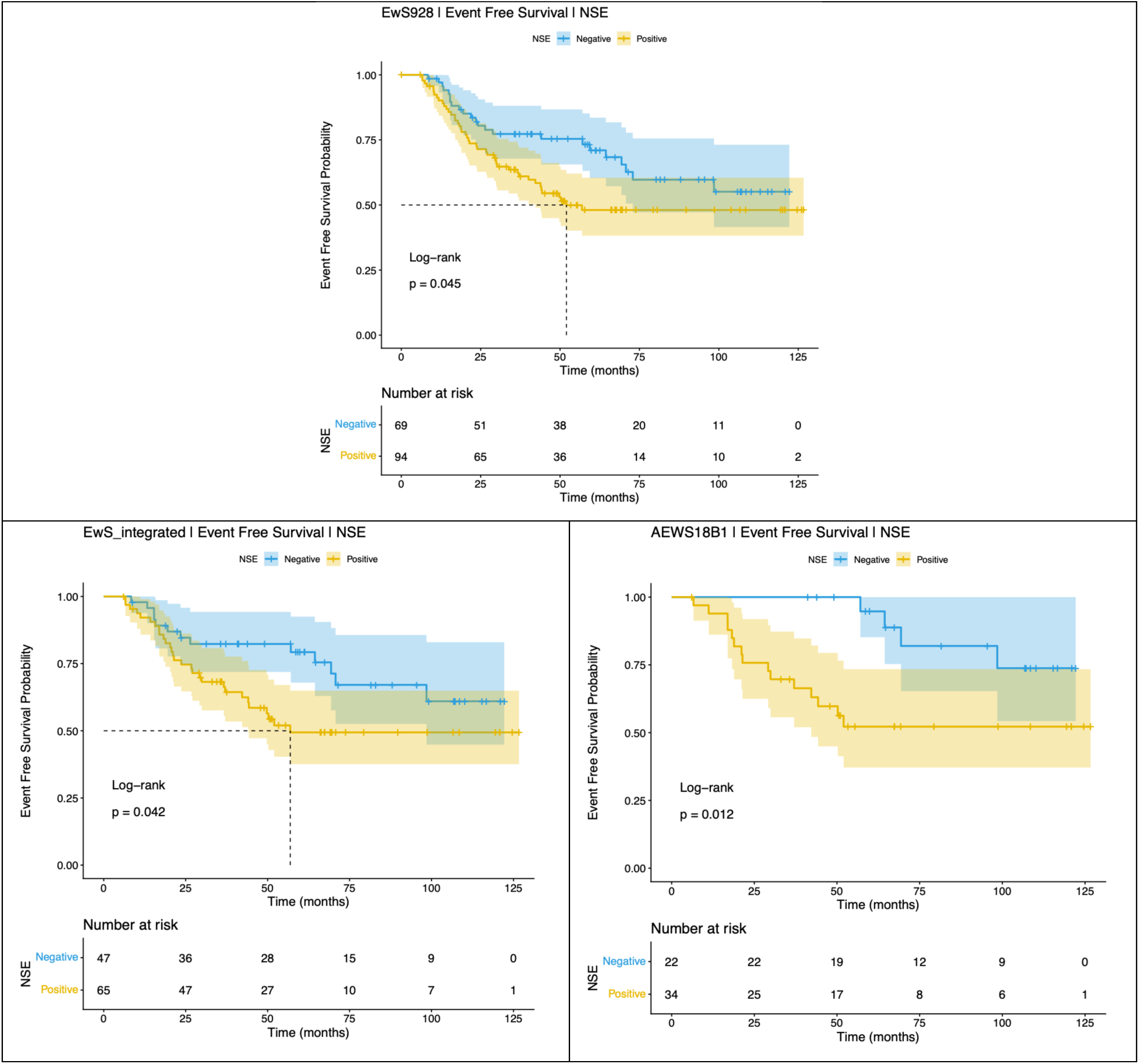
Kaplan–Meier curves for event-free survival by NSE status in the three analysis sets.

**Figure S4.**
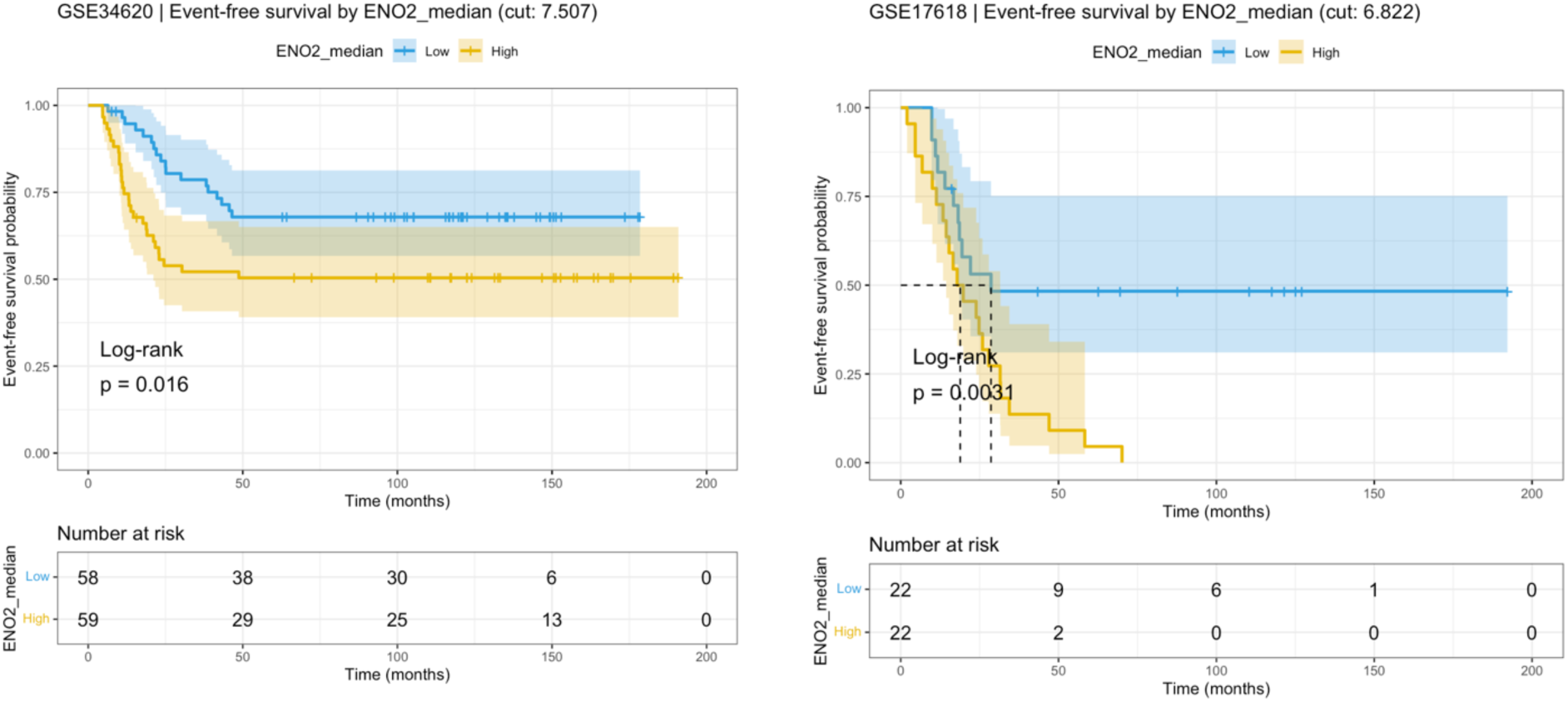
Kaplan–Meier curves for EFS by *ENO2* expression in GSE34620 and GSE17618.

**Figure S5.**
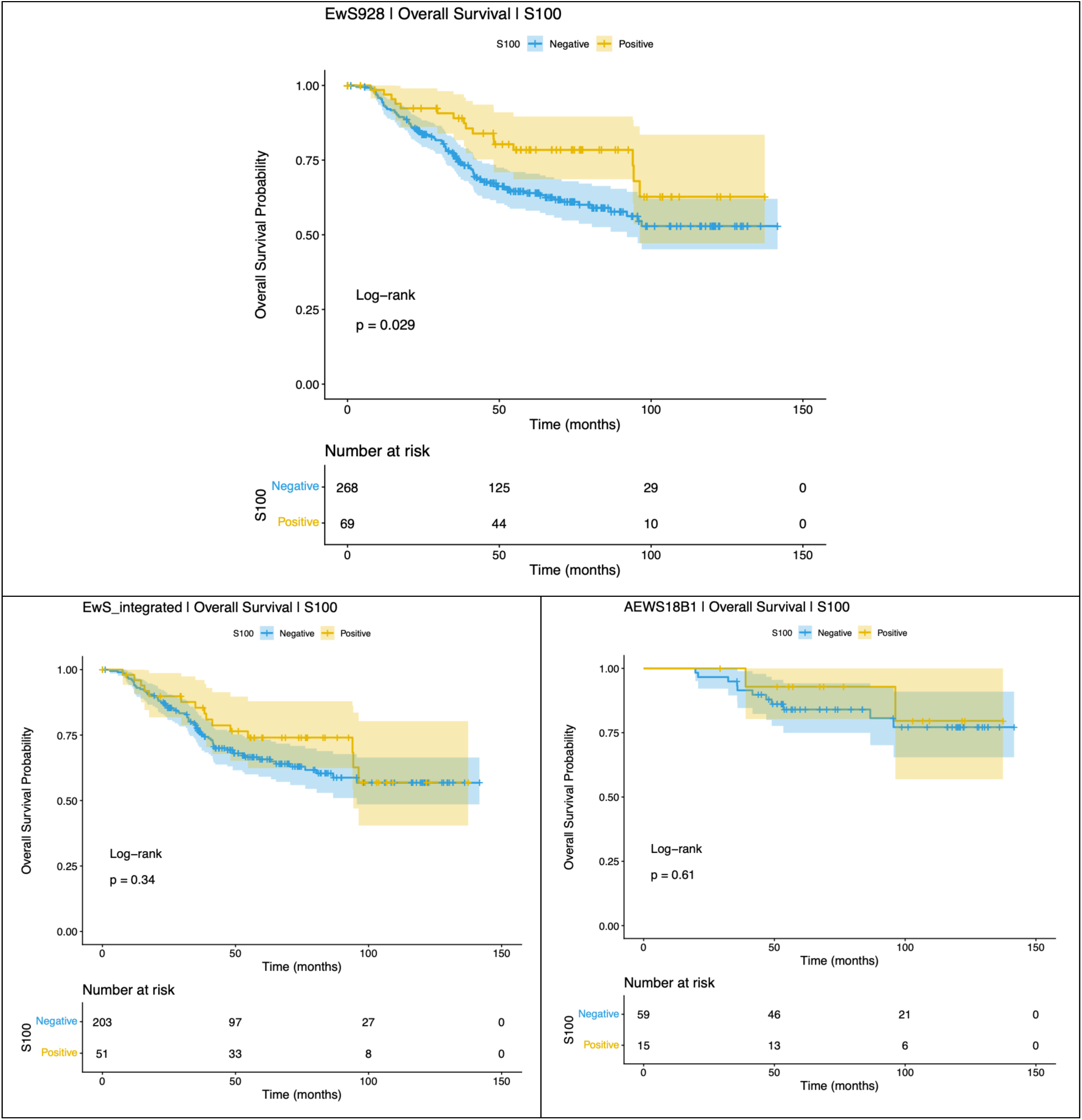
Kaplan–Meier curves for OS by S100 status in the full cohort and the two nested molecularly defined subsets.

**Table S1.** Survival by NSE and S100 status in each analysis population (A), and adjusted, subgroup and interaction models for NSE (B), including event counts and Firth penalized estimates.

| **Marker** | **Tier** | **Endpoint** | **Negative n (events)** | **Positive n (events)** | **5-year survival % (95% CI) Negative** | **5-year survival % (95% CI) Positive** | **HR (95% CI)** | **Log-rank p** |
|---|---|---|---|---|---|---|---|---|
| NSE | Discovery | OS | 69 (14) | 93 (35) | 84.6 (76.3-93.9) | 60.5 (50.4-72.7) | 2.17 (1.17-4.05) | 0.012 |
| NSE | Discovery | EFS | 69 (23) | 94 (44) | 71.0 (60.4-83.6) | 48.1 (38.2-60.5) | 1.67 (1.01-2.77) | 0.045 |
| NSE | Molecularly confirmed | OS | 47 (9) | 65 (26) | 84.7 (75.0-95.8) | 59.0 (47.1-74.0) | 2.46 (1.15-5.27) | 0.017 |
| NSE | Molecularly confirmed | EFS | 47 (13) | 65 (29) | 79.3 (67.9-92.5) | 49.4 (37.6-64.9) | 1.95 (1.01-3.77) | 0.042 |
| NSE | Fusion confirmed | OS | 22 (0) | 34 (11) | 100 (80.5-100) | 68.7 (53.2-88.9) | 20.1 (2.60-2586) | 0.002 (Firth<sup>*</sup>) |
| NSE | Fusion confirmed | EFS | 22 (4) | 34 (15) | 94.7 (85.2-100) | 52.2 (37.1-73.4) | 3.80 (1.25-11.6) | 0.012 |
| S100 | Discovery | OS | 268 (97) | 69 (16) | 64.0 (58.1-70.5) | 78.4 (68.6-89.6) | 0.56 (0.33-0.95) | 0.029 |
| S100 | Discovery | EFS | 268 (115) | 69 (22) | 54.8 (48.9-61.5) | 66.6 (55.4-80.0) | 0.65 (0.41-1.03) | 0.067 |
| S100 | Molecularly confirmed | OS | 203 (69) | 51 (15) | 65.8 (59.1-73.3) | 74.1 (62.4-87.9) | 0.76 (0.44-1.33) | 0.34 |
| S100 | Molecularly confirmed | EFS | 203 (86) | 51 (18) | 55.5 (48.8-63.2) | 62.1 (49.1-78.5) | 0.76 (0.46-1.27) | 0.3 |
| S100 | Fusion confirmed | OS | 59 (11) | 15 (2) | 84.0 (74.9-94.2) | 92.9 (80.3-100) | 0.67 (0.15-3.04) | 0.61 |
| S100 | Fusion confirmed | EFS | 59 (17) | 15 (5) | 72.7 (62.1-85.1) | 68.9 (47.5-100) | 1.05 (0.39-2.85) | 0.92 |

**Table S2.**
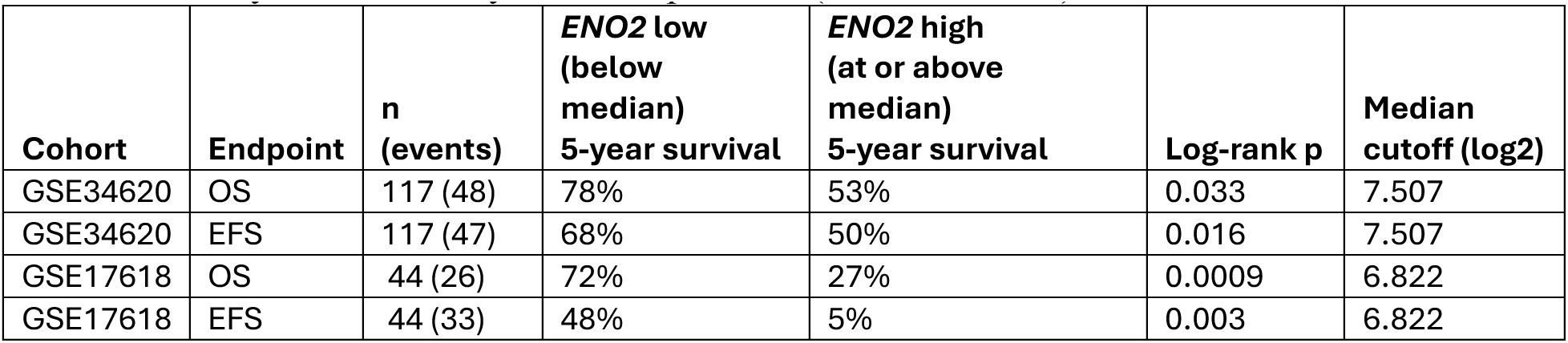
Five-year survival by *ENO2* expression (median cut-off) in the external cohorts.

**Table S3.** Univariable screen of all evaluable immunostains for EFS across the three nested populations, with Benjamini–Hochberg adjustment within each population.

| Tier | End point | Marker | Negative n (events) | Positive n (events) | HR (95% CI) | Log-rank p | BH-adjusted p | Markers tested |
|---|---|---|---|---|---|---|---|---|
| Discovery | OS | NSE | 69 (14) | 93 (35) | 2.17 (1.17-4.05) | 0.012 | 0.12 | 11 |
| Discovery | OS | S100 | 268 (97) | 69 (16) | 0.56 (0.33-0.95) | 0.029 | 0.12 | 11 |
| Discovery | OS | Desmin | 576 (180) | 8 (4) | 2.80 (1.04-7.55) | 0.034 | 0.12 | 11 |
| Discovery | OS | LCA (CD45) | 574 (174) | 5 (3) | 2.16 (0.69-6.78) | 0.17 | 0.48 | 11 |
| Discovery | OS | FLI-1 | 19 (8) | 201 (60) | 0.68 (0.32-1.42) | 0.30 | 0.66 | 11 |
| Discovery | OS | Chromogranin | 171 (49) | 7 (1) | 0.47 (0.06-3.37) | 0.44 | 0.71 | 11 |
| Discovery | OS | MyoD1 | 117 (36) | 6 (1) | 0.47 (0.06-3.45) | 0.45 | 0.71 | 11 |
| Discovery | OS | ERG | 20 (3) | 17 (4) | 1.55 (0.34-6.96) | 0.57 | 0.78 | 11 |
| Discovery | OS | Vimentin | 13 (4) | 191 (52) | 0.80 (0.29-2.22) | 0.67 | 0.82 | 11 |
| Discovery | OS | WT1 | 184 (64) | 8 (3) | 1.19 (0.37-3.78) | 0.77 | 0.85 | 11 |
| Discovery | OS | Synaptophysin | 228 (73) | 154 (48) | 1.00 (0.70-1.44) | 1.00 | 1.00 | 11 |
| Discovery | EFS | NSE | 69 (23) | 94 (44) | 1.67 (1.01-2.77) | 0.045 | 0.37 | 11 |
| Discovery | EFS | S100 | 268 (115) | 69 (22) | 0.65 (0.41-1.03) | 0.067 | 0.37 | 11 |
| Discovery | EFS | Desmin | 578 (237) | 8 (4) | 1.88 (0.70-5.07) | 0.20 | 0.72 | 11 |
| Discovery | EFS | FLI-1 | 19 (9) | 201 (70) | 0.67 (0.34-1.35) | 0.26 | 0.72 | 11 |
| Discovery | EFS | WT1 | 183 (76) | 8 (4) | 1.20 (0.44-3.28) | 0.72 | 0.98 | 11 |
| Discovery | EFS | MyoD1 | 118 (52) | 6 (3) | 1.18 (0.37-3.78) | 0.78 | 0.98 | 11 |
| Discovery | EFS | Synaptophysin | 228 (96) | 154 (64) | 1.03 (0.75-1.42) | 0.84 | 0.98 | 11 |
| Discovery | EFS | Vimentin | 13 (5) | 193 (82) | 1.10 (0.44-2.71) | 0.84 | 0.98 | 11 |
| Discovery | EFS | ERG | 20 (5) | 17 (4) | 1.10 (0.29-4.08) | 0.89 | 0.98 | 11 |
| Discovery | EFS | LCA (CD45) | 575 (232) | 5 (2) | 1.07 (0.27-4.30) | 0.93 | 0.98 | 11 |
| Discovery | EFS | Chromogranin | 171 (68) | 7 (3) | 0.98 (0.31-3.12) | 0.98 | 0.98 | 11 |
| Molecularly confirmed | OS | NSE | 47 (9) | 65 (26) | 2.46 (1.15-5.27) | 0.017 | 0.13 | 8 |
| Molecularly confirmed | OS | LCA (CD45) | 439 (132) | 5 (3) | 2.25 (0.72-7.08) | 0.15 | 0.61 | 8 |
| Molecularly confirmed | OS | S100 | 203 (69) | 51 (15) | 0.76 (0.44-1.33) | 0.34 | 0.68 | 8 |
| Molecularly confirmed | OS | ERG | 15 (2) | 13 (3) | 1.98 (0.33-11.84) | 0.45 | 0.68 | 8 |
| Molecularly confirmed | OS | WT1 | 142 (47) | 5 (1) | 0.57 (0.08-4.15) | 0.57 | 0.68 | 8 |
| Molecularly confirmed | OS | FLI-1 | 13 (5) | 163 (47) | 0.77 (0.31-1.93) | 0.58 | 0.68 | 8 |
| Molecularly confirmed | OS | Vimentin | 10 (2) | 140 (40) | 1.47 (0.36-6.10) | 0.59 | 0.68 | 8 |
| Molecularly confirmed | OS | Synaptophysin | 169 (55) | 119 (36) | 0.92 (0.61-1.41) | 0.71 | 0.71 | 8 |
| Molecularly confirmed | EFS | NSE | 47 (13) | 65 (29) | 1.95 (1.01-3.77) | 0.042 | 0.34 | 8 |
| Molecularly confirmed | EFS | S100 | 203 (86) | 51 (18) | 0.76 (0.46-1.27) | 0.30 | 0.89 | 8 |
| Molecularly confirmed | EFS | Vimentin | 10 (3) | 141 (60) | 1.59 (0.50-5.07) | 0.43 | 0.89 | 8 |
| Molecularly confirmed | EFS | FLI-1 | 13 (5) | 163 (53) | 0.77 (0.31-1.93) | 0.58 | 0.89 | 8 |
| Molecularly confirmed | EFS | Synaptophysin | 169 (67) | 119 (48) | 1.05 (0.73-1.52) | 0.79 | 0.89 | 8 |
| Molecularly confirmed | EFS | ERG | 15 (4) | 13 (3) | 1.17 (0.26-5.24) | 0.84 | 0.89 | 8 |
| Molecularly confirmed | EFS | LCA (CD45) | 440 (168) | 5 (2) | 1.14 (0.28-4.61) | 0.85 | 0.89 | 8 |
| Molecularly confirmed | EFS | WT1 | 142 (56) | 5 (2) | 0.90 (0.22-3.70) | 0.89 | 0.89 | 8 |
| Fusion confirmed | OS | NSE | 22 (0) | 34 (11) | 20.11 (2.60-2586)† | 0.002 | 0.008 | 4 |
| Fusion confirmed | OS | FLI-1 | 5 (2) | 55 (11) | 0.55 (0.12-2.47) | 0.42 | 0.61 | 4 |
| Fusion confirmed | OS | Synaptophysin | 55 (9) | 39 (8) | 1.39 (0.54-3.62) | 0.49 | 0.61 | 4 |
| Fusion confirmed | OS | S100 | 59 (11) | 15 (2) | 0.67 (0.15-3.04) | 0.61 | 0.61 | 4 |
| Fusion confirmed | EFS | NSE | 22 (4) | 34 (15) | 3.80 (1.25-11.56) | 0.012 | 0.046 | 4 |
| Fusion confirmed | EFS | Synaptophysin | 55 (14) | 39 (13) | 1.46 (0.69-3.11) | 0.32 | 0.64 | 4 |
| Fusion confirmed | EFS | FLI-1 | 5 (2) | 55 (16) | 0.74 (0.17-3.24) | 0.70 | 0.92 | 4 |
| Fusion confirmed | EFS | S100 | 59 (17) | 15 (5) | 1.05 (0.39-2.85) | 0.92 | 0.92 | 4 |

Notes: A marker is evaluable only if both the negative and the positive group contain at least 5 patients.

**Table S4.** *ENO2* expression and survival by tertile in the external cohorts.

| Cohort | Endpoint | *ENO2* tertile | Tertile boundaries (log2) | n (events) | 5-year survival % (95% CI) | HR vs Low (95% CI) | Wald p | Log-rank p (3 groups) |
|---|---|---|---|---|---|---|---|---|
| GSE34620 | OS | Low | 6.689 / 8.052 | 39 (13) | 79 (68-93) | 1 (reference) |  | 0.062 |
| GSE34620 | OS | Mid | 6.689 / 8.052 | 39 (14) | 67 (53-83) | 1.22 (0.57-2.59) | 0.61 | 0.062 |
| GSE34620 | OS | High | 6.689 / 8.052 | 39 (21) | 49 (35-67) | 2.14 (1.07-4.27) | 0.032 | 0.062 |
| GSE34620 | EFS | Low | 6.689 / 8.052 | 39 (12) | 68 (55-85) | 1 (reference) |  | 0.048 |
| GSE34620 | EFS | Mid | 6.689 / 8.052 | 39 (15) | 61 (47-78) | 1.40 (0.66-3.00) | 0.38 | 0.048 |
| GSE34620 | EFS | High | 6.689 / 8.052 | 39 (20) | 48 (35-67) | 2.34 (1.14-4.80) | 0.02 | 0.048 |
| GSE17618 | OS | Low | 6.467 / 7.352 | 15 (6) | 67 (47-95) | 1 (reference) |  | 0.028 |
| GSE17618 | OS | Mid | 6.467 / 7.352 | 14 (7) | 62 (40-95) | 1.35 (0.45-4.03) | 0.59 | 0.028 |
| GSE17618 | OS | High | 6.467 / 7.352 | 15 (13) | 20 (7-55) | 3.20 (1.20-8.52) | 0.02 | 0.028 |
| GSE17618 | EFS | Low | 6.467 / 7.352 | 15 (9) | 40 (22-74) | 1 (reference) |  | 0.068 |
| GSE17618 | EFS | Mid | 6.467 / 7.352 | 14 (9) | 29 (12-71) | 1.10 (0.43-2.78) | 0.84 | 0.068 |
| GSE17618 | EFS | High | 6.467 / 7.352 | 15 (15) | 7 (1-44) | 2.32 (1.00-5.37) | 0.049 | 0.068 |

Notes: *ENO2* tertiles defined within each cohort. Hazard ratios are from a univariable Cox model with the lowest tertile as reference; the log-rank p compares all three groups. 5-year survival is Kaplan-Meier with log-transformed confidence intervals. In GSE17618 the tertile groups hold 14 to 15 patients, so the estimates are imprecise.

**Table S5.** ENO2 (high vs low) and survival in the external cohorts over all follow-up and by follow-up period, with proportional-hazards diagnostics.

| Cohort | Endpoint | Period | Patients at risk (events) | HR (95% CI) | Wald p | ENO2 x period interaction p | Schoenfeld p |
|---|---|---|---|---|---|---|---|
| GSE34620 | OS | All follow-up | 117 (48) | 1.86 (1.04-3.32) | 0.036 |  | 0.063 |
| GSE34620 | OS | 0-24 months | 117 (24) | 3.23 (1.28-8.14) | 0.013 | 0.1 |  |
| GSE34620 | OS | >24 months | 93 (24) | 1.16 (0.52-2.60) | 0.71 | 0.1 |  |
| GSE34620 | EFS | All follow-up | 117 (47) | 2.04 (1.13-3.67) | 0.018 |  | 0.002 |
| GSE34620 | EFS | 0-24 months | 117 (35) | 3.53 (1.65-7.53) | 0.001 | 0.009 |  |
| GSE34620 | EFS | >24 months | 79 (12) | 0.47 (0.13-1.73) | 0.26 | 0.009 |  |
| GSE17618 | OS | All follow-up | 44 (26) | 3.94 (1.65-9.43) | 0.002 |  | 0.4 |
| GSE17618 | OS | 0-24 months | 44 (14) | 2.93 (0.92-9.36) | 0.069 | 0.49 |  |
| GSE17618 | OS | >24 months | 29 (12) | 5.46 (1.47-20.3) | 0.011 | 0.49 |  |
| GSE17618 | EFS | All follow-up | 44 (33) | 2.91 (1.39-6.11) | 0.005 |  | 0.065 |
| GSE17618 | EFS | 0-24 months | 44 (23) | 1.52 (0.67-3.47) | 0.32 | 0.018 |  |
| GSE17618 | EFS | >24 months | 20 (10) | 23.1 (2.82-189) | 0.003 | 0.018 |  |

Notes:
1. Hazard ratios (HR) with 95% confidence intervals (CI) are for ENO2 high vs low, defined by the within-cohort median expression (Methods). All models are unadjusted.
2. All follow-up: the univariable Cox model over the entire follow-up, as reported in the main text. Schoenfeld p is the proportional-hazards test for this model (cox.zph); a value below 0.05 indicates that the ENO2 hazard ratio is not constant over follow-up.
3. 0-24 months and >24 months: a single Cox model per cohort and endpoint in which follow-up was split at 24 months and ENO2 was given a separate coefficient in each period, equivalent to a time-dependent covariate model with a step at 24 months. The ENO2 x period interaction p (Wald) tests whether the two period HRs differ and is shown once per model.
4. Patients at risk (events): the number of patients entering the period and, in parentheses, the events occurring within it.
5. Beyond 24 months the GSE17618 estimates rest on 10 to 12 events and are imprecise.

